# Estimation of Full-Length TprK Diversity in *Treponema pallidum* subspecies *pallidum*

**DOI:** 10.1101/2020.01.23.917484

**Authors:** Amin Addetia, Michelle Lin, Quynh Phung, Hong Xie, Meei-Li Huang, Giulia Ciccarese, Ivano Dal Conte, Marco Cusini, Francesco Drago, Lorenzo Giacani, Alexander L. Greninger

**Author notes:** Address correspondence to Alexander L. Greninger.

## Abstract

Immune evasion and disease progression of *Treponema pallidum* subspecies *pallidum* are associated with sequence diversity in the hypervariable, putative outer membrane protein TprK. Previous attempts to study variation within TprK have sequenced at depths insufficient to fully appreciate the hypervariable nature of the protein, failed to establish linkage between the protein’s 7 variable regions, or were conducted on strains passed through rabbits. As a consequence, a complete profiling of *tprK* during infection in the human host is still lacking. Furthermore, prior studies examining how *T. pallidum* uses its repertoire of genomic donor sites to generate diversity within the V regions of the *tprK* also yielded a partial understanding of this process, due to the limited number of *tprK* alleles examined. In this study, we used short- and long-read deep sequencing to directly characterize full-length *tprK* alleles from *T. pallidum* collected from early lesions of patients attending two STD clinics in Italy. Our data, combined with recent data available on Chinese *T. pallidum* strains, show the near complete absence of overlap in TprK sequences among the 41 strains profiled to date. Moreover, our data allowed us to redefine the boundaries of *tprK* V regions, identify 55 donor sites, and estimate the total number of TprK variants that *T. pallidum* can potentially generate. Altogether, our results support how *T. pallidum* TprK antigenic variation system is an unsurmountable obstacle for the human immune system to naturally achieve infection eradication, and reiterate the importance of this mechanism for pathogen persistence in the host.

**Importance:** Syphilis continues to be a significant public health issue in both low- and high-income nations, including the United States, where the number of infectious syphilis cases has increased dramatically over the past five years. *T. pallidum*, the causative agent of syphilis, encodes an outer membrane protein TprK that undergoes segmental gene conversion to constantly create new sequences. We performed deep TprK profiling to understand full-length TprK diversity in *T. pallidum-*positive clinical specimens and compared these to all samples for which TprK deep sequencing is available. We found almost no overlap in TprK sequences between different patients. We further estimate that the total baseline junctional diversity of full-length TprK rivals that of current estimates of the human adaptive immune system. These data underscore the immunoevasive ability of TprK that allows *T. pallidum* to establish lifelong infection.

## Introduction

Syphilis, caused by the spirochete *Treponema pallidum* subspecies *pallidum* (*T. pallidum*), is a significant global health problem. Although most syphilis cases occur in low-income countries, where the disease is endemic, rates of syphilis infection have been steadily increasing for the last two decades in high-income nations, particularly in men who have sex with men (MSM) and HIV-infected individuals [1–4]. Syphilis is an acute and chronic sexually transmitted infection marked by distinct early and late stages [5]. These stages are generally distinguished by unique clinical manifestations with symptoms associated with the late stage developing up to several decades after initial infection and following a long period of disease latency [6].

The mechanisms that allow *T. pallidum* to persist for the lifetime of an infected individual are not fully understood. During natural and experimental syphilis infection, a robust host immune response is developed against *T. pallidum* [7]. This suggests immune evasion strategies developed by *T. pallidum* are a key aspect of its pathogenesis [8].

The ability of *T. pallidum* to evade the host immune response is attributed to the organism’s scarcity of surface-exposed outer membrane proteins (OMPs), very slow generation time (∼33h), and the ability to stochastically and rapidly switch on and off expression of genes encoding putative OMPs through phase variation [9]. Chief among the immune evasion strategies evolved by *T. pallidum* is its ability to generate diversity within the putative OMP TprK [10–12]. TprK harbors seven discrete variable (V) regions, V1-V7. In the putative TprK beta-barrel structure, each variable region is predicted to form a loop exposed at the host-pathogen interface [13]. Generation of variants in these V regions occurs through non-reciprocal segmental gene conversion, a process in which sections from donor sites flanking the *tprD* gene (*tp0117*) are stitched together to create new sequences [14,15]. Forty-seven putative donor sites have been identified thus far [15], however, the total number of unique TprK sequences that can be generated in a *T. pallidum* strain has yet to be determined.

Gene conversion results in significant intra- and inter-strain diversity of the TprK protein [16,17,14,18,19]. In rabbit models, diversity in TprK actively accumulates over the course of an infection and appears to be driven by the host’s immune response [20,21]. At least five of the variable regions, V2 and V4-V7, of TprK elicit an antibody response in rabbit models [22]. These antibodies are specific for a single variable sequence [22], which further supports that generation of new V region sequences allows *T. pallidum* to evade the host response. Furthermore, increased diversity of TprK is directly correlated with more advanced stages of syphilis [19,23]. In both rabbit models and humans, *T. pallidum* strains isolated from cases of secondary/disseminated syphilis contained more TprK diversity than those isolated from cases of primary syphilis [19,23].

Previous studies to evaluate TprK variability within *T. pallidum* strains have sequenced of a limited number of TprK clones, failed to resolve linkage between variable regions, or been conducted on strains passed through rabbits [16,18–20] [24]. As a result, no studies to date have adequately profiled TprK within *T. pallidum* during natural infection in the human host. Furthermore, understanding of how different donor sites contribute to variable region sequences has been hindered by the analysis using a limited number of *tprK* clones [25]. In this study, we used short- and long-read deep sequencing to directly characterize full-length TprK genes amplified from *T. pallidum* collected from early genital or anal lesions of 13 individuals attending two STD clinics in Milan and Turin, Italy [26]. We then combine our data with recent short-read *tprK* sequencing data from 28 *T. pallidum* strains from collected in China to illustrate the near complete lack of overlap in TprK sequences among all 41 clinical strains directly and deeply profiled to date. Additionally, our data help to redefine the TprK variable regions and to provide an estimate of the number of TprK variants that *T. pallidum* can potentially generate with its repertoire of donor cassettes. Overall, our data reiterate the pivotal importance of the TprK antigenic variation system to allow *T. pallidum* persistence in the host during infection.

## Methods

### Sample collection

Swabs from genital or anal lesions were collected from syphilis patients attending the Dermatology Clinics of the University of Turin and the Ospedale Maggiore in Milan from approximately December 2016 to March 2017. The only exclusion criterion for sample collection was an existing record of antibiotic therapy initiated within 30 days from the patient visit. For sample collection, whenever possible, the lesion area was gently squeezed to imbibe the swabs with exudate. The swabs were then placed in sterile microcentrifuge tubes containing 1 ml of 1X lysis buffer (10 mM Tris-HCl, 0.1 M ethylenediaminetetraacetic acid, and 0.5% sodium dodecyl-sulfate) suitable for DNA extraction. The swab shafts were then cut to leave the swab in the buffer. Samples were kept frozen at −80°C until DNA extraction. Sample collection was authorized by the Human Subject Committee of each collecting institution (Protocol code: PR033REG2016 for the University of Turin, and Protocol Code TREPO2016 for the University of Milan) and informed consent was obtained from each patient. Specimens were then sent as de-identified samples in dry ice to the University of Washington for DNA extraction. The University of Washington Institutional Review Board determined this investigation not to be human subject research. Patient demographics were also collected as well as information on sexual orientation, HIV status, syphilis stage and serology results (VDRL/RPR and TPHA/TPPA) at the time of patient visit.

### DNA extraction and strain typing

Frozen samples were thawed at room temperature and vortexed before processing. DNA was extracted from 200 µl of sample suspension using the QIAamp DNA mini kit (Qiagen, Valencia, CA) according to the manufacturer’s instructions. DNA was resuspended in 100 µl of elution buffer provided with the kit. Successful DNA extraction was checked by amplification of a fragment of the human β-globin gene (Sense primer 5′-CAA CTT CAT CCA CGT TCA CC-3′, Antisense primer 5′-GAA GAG CCA AGG ACA GGT A-3′; expected size: 268 bp). Amplifications were performed in a 50 µl final volume using 5 µl of DNA template and 2.5 units of GoTaq polymerase (Promega, Madison, WI). Final concentrations of MgCl_2_, dNTPs, and each primer were 1.5 mM, 200 µM, and 0.32 µM, respectively. Cycling conditions were initial denaturation at 95°C for 4 minutes, followed 95°C for 1 min, 60°C for 1 min and 72°C for 1 min for a total of 40 cycles. Final extension was at 72°C for 5 min.

### Quantification of Treponemal Load within Patient Samples

The treponemal load of each sample was measured by qPCR as previously described [24]. Briefly, a portion of *tp47* was amplified using 14.33 µL of 2x QuantiTect multiplex PCR mix, 0.65 µL of 2x QuantiTect multiplex PCR mix with ROX, 0.03 unit of UNG and the following primers 5′-CAA GTA CGA GGG GAA CAT CGA T-3′ and 5′-TGA TCG CTG ACA AGC TTA GG-3′. Amplification was monitored with the following probe: 5′-FAM-CGG AGA CTC TGA TGG ATG CTG CAG TT-NFQMGB-3′. The following conditions were used for the qPCR reaction: 50°C for 2 minutes, 95°C for 15 minutes, and 45 cycles of 94°C for 1 minute and 60°C for 1 minute.

### Direct from sample amplification and next-generation sequencing of tprK

PCR amplification of *tprK* was conducted using previously described conditions [24] and *tprK*-specific primers appended to 16 bp Pacbio barcodes (Table S1). The resulting 1.7kb product was cleaned using 0.6x volumes of AMPure XP beads (Beckman-Coulter). For long-read sequencing, library construction and sequencing on a Sequel I SMRT Cell 1M with a 10-hour movie were completed by the University of Washington PacBio Sequencing Services. A minimum of 5,224 PacBio reads were obtained for each of the samples. Short-read libraries from the same full-length amplicons were constructed with the Nextera XT kit (Illumina), cleaned with 0.6x volumes of AMPure XP beads (Beckman-Coulter), and sequenced on 1×192 bp Illumina MiSeq runs. A minimum of 101,000 Illumina sequencing reads, corresponding to a minimum mean coverage of 6,672x, were obtained for each sample. Sequencing metadata is available in Table S2.

### Sequencing analysis

Analysis of *tprK* was performed using custom python/R scripts available on GitHub (https://github.com/greninger-lab/tprK_diversity). For the Italian samples, because of the tagmentation-based library preparation, we quality- (Q20) and adapter-trimmed Illumina reads using trimmomatic v0.38. PacBio Q20 CCS reads between 1400-1800 bp were trimmed of PCR primers using the dada2 preprocessing pipeline and denoised using RAD [27,28]. Previously published short-read tiling sequencing data for *tprK* from 14 primary and 14 secondary syphilis infections in adults from Xiamen University was downloaded from the NCBI Sequence Read Archive [29,30]. Because of the tiling PCR library design followed by 2×300bp, both paired-end reads were used in analysis of the Xiamen samples after adapter trimming using the same options as above. Variable regions were extracted from all samples using fuzzy regular expression matching using 18bp of neighboring conserved sequence with up to a 3bp mismatch. Because of the slight differences in coverage, we required a minimum of 5 reads of support for a given variable region amino acid sequence from the Xiamen samples, while for the Italian samples we used a minimum of 10 reads. We additionally included short-read sequencing data from 2 *T. pallidum* strains passaged in rabbits, which we profiled in a previous investigation [24], in our analysis. Similar to the Italian strains, we required each unique identified variable region sequence in these strains to be supported by a minimum of 10 sequencing reads and present at or above a frequency of 0.2%.

For full-length TprK phylogenetic analysis, we removed any TprK sequences that contained stop codons or which failed to fuzzy match a 20 amino acid region (allowing 3 mismatches) in any conserved region abutting a variable region, which we found was indicative of two frame shifts in consecutive variable regions in two TprK sequences. We also removed any full-length TprK sequences that comprised <0.2% of sequences present in a given sample for the purposes of display.

We used blastn with exact matching over a word size of 10 – our estimate of the smallest, high-confidence contribution of a donor site – to identify potential donor sites within a 17.2kb locus containing the *tprD* gene. We limited the number of potential contributions of each donor site to a variable region to three by restricting the maximum high scoring pairs (-max_hsps 3). We used the subject_besthit option to force non-overlapping HSPs. In order to generate a list of high-confidence donor sites and reduce putative false positives due to the smaller word size and to control for potential sequencing error, we only used variable regions with greater than 50 reads of support and 0.2% frequency (within-sample) and also required donor sites to be used in recovered *tprK* variable region sequences in at least 2 separate samples.

Shannon diversity scores for each sample were calculated using the R package VEGAN [31]. Differences in the number of variable region sequences and diversity scores for strains stratified by host factors were assessed using the Wilcoxon Rank-Sum test.

### Data availability

Illumina and PacBio reads from *tprK* sequencing of the samples used in this study are available under the NCBI BioProject number PRJNA589065.

## Results

### Italian patient metadata

We selected 13 *T. pallidum* strains collected from syphilis patients, comprising 7 primary and 6 secondary syphilis cases, in Milan and Turin, Italy (Table 1, Table S3). All patients reported to be MSM and the median age of individuals was 39 years (range 20-57 years). Eight of the individuals sampled were HIV positive and, for nine of the patients, this was the first syphilis diagnosis. Seven of the specimens were collected from genital lesions, while the remaining six were collected from anal lesions.

**Table 1.**
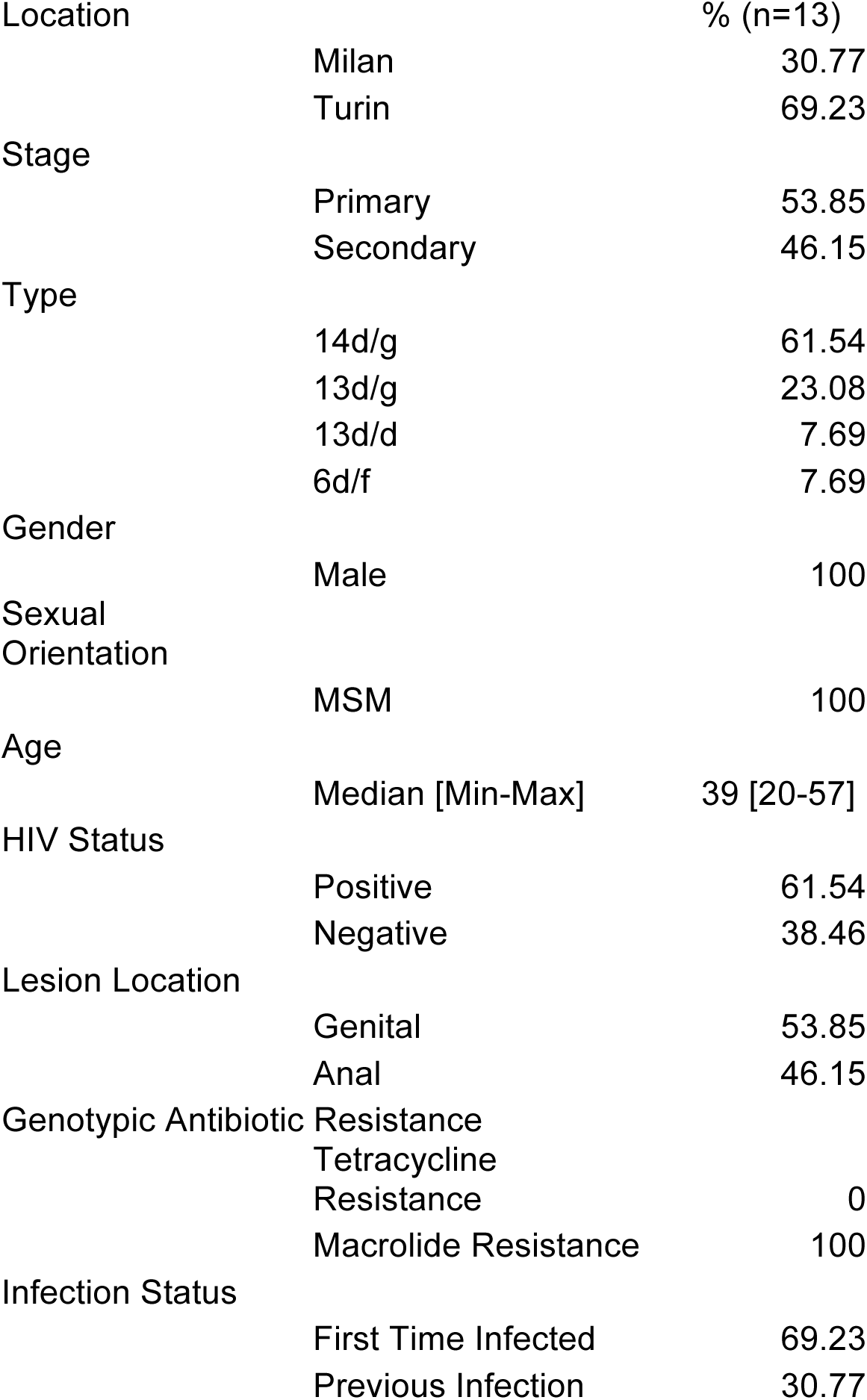
Summary statistics of patient metadata for strains sequenced in this study

### T. pallidum DNA quantitation from clinical specimens

We first assessed the impact of the amount of treponemal DNA input into our initial PCR reaction on our ability to detect diversity within the 7 variable regions of *tprK*. For 3 strains, we compared inputs of 1,000 copies of treponemal DNA to the maximum possible input for our *tprK* PCR amplification reaction. Additionally, we performed a technical replicate using the same copy number input for another strain. For strains AS10, AS11, and AS12, the maximum input for *tprK* PCR was 5,362, 2,736, 6,663 copies of treponemal DNA. For strain AS18, we repeated the *tprK* PCR with 1,013 copies of treponemal DNA. The number of identified variants and the diversity measures for each variable region were similar despite the varying inputs (Table S4, Figure S1). For our subsequent analyses, we normalized the input for the initial *tprK* amplification to 1,000 treponemal copies for each sample.

### TprK diversity in T. pallidum strains directly sampled from humans

We used short-read sequencing to examine the diversity within the seven V regions of TprK and required each identified amino acid sequence from an isolate to be supported by a minimum of 10 sequencing reads and present at a relative frequency greater than or equal to 0.2%. We identified a median of 65 (range: 37-162) unique sequences from all seven V regions from our 13 *T. pallidum* strains. Across the 13 strains, V1 contained the overall fewest unique sequences (median: 4, range: 1-7) and, as determined by the Shannon diversity index, was the least diverse variable region (median: 0.119, range: 0-0.836). V6 contained the greatest number of unique variants (median: 20; range: 3-65) and was also the most diverse variable region (median: 1.603; range: 0.363-2.760) (Table S5).

We next examined the diversity of TprK in the context of different clinical characteristics. *T. pallidum* strains collected from cases of secondary syphilis contained significantly more unique variable region sequences (p=0.004) and were significantly more diverse (p=0.002) than those strains collected from cases of primary syphilis (Table 2). The number of unique sequences did not significantly differ (p=0.174) between strains collected from anal or genital lesions. However, strains collected from anal lesions exhibited significantly more diversity (p=0.035) across the seven V regions. No significant differences were observed in the number of unique variants or diversity when stratified by HIV status of the patient (p=0.187; p=0.171) or history of prior *T. pallidum* infection (p=0.537; p=0.711).

**Table 2.**
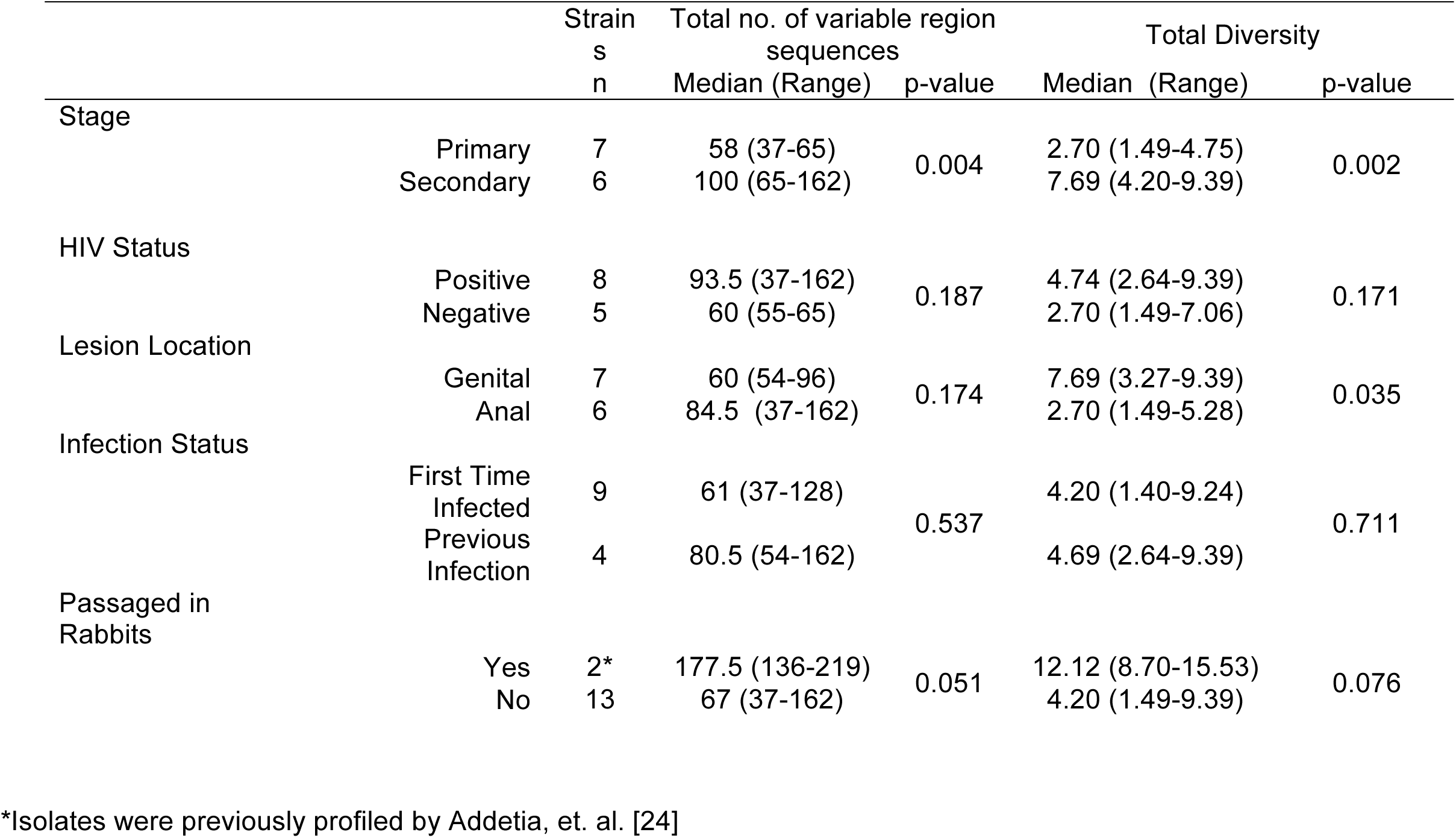
Comparison of the total number of variable region sequences and total Shannon diversity scores across the 7 variable regions of TprK in the context of clinical characteristics and passage history.

In a previous investigation, we profiled TprK in two *T. pallidum* strains collected from a single patient and amplified by two passages of strains in New Zealand white rabbits [24]. To assess the impact of the additional passage through rabbits on TprK, we compared the number of unique variants and diversity across the seven V regions identified from the 13 Italian *T. pallidum* strains and our two previously profiled strains. Strains passed through rabbits contained a greater number of variable region sequences (median 177.5 vs. 65, p=0.051) and greater diversity across the seven variable regions (median 12.1 vs. 4.2, p=0.076) compared to those directly sequences from clinical samples, though these differences were not significant given the few rabbit strains previously sampled.

To ensure accurate estimation of variable regions in both unlinked and linked analyses, we compared results from both short-read (Illumina) and long-read (PacBio) sequencing of all *tprK* amplicons generated in this study. The variable region allele percentages as measured by short-read and long-read sequencing were highly correlated (median r^2^ = 0.995, range 0.974-1.00) (Figure S2), illustrating the high quality of modern long-read sequencing. PacBio sequencing exhibited an overall positive bias in variable region allele percentage compared to Illumina sequencing with an average linear regression slope of 1.029 (range 1.011-1.092), likely due to clustering during read denoising and post-filtering.

Using our long read data, we recovered a total of 634 full-length TprKs across the 13 samples, ranging from 26-95 different full-length TprKs within each sample at ≥ 0.2% frequency. The most prevalent TprK in each sample was generally located near the root of the TprK phylogenetic tree for that particular sample (Figure 1). We found that only 3 full-length TprK sequences were shared among the 634 TprK sequences recovered from all 13 patients after removing sequences at a frequency of <0.2% within each sample. Two of these overlapping TprK sequences comprised the most common sequences in at least one of the samples. For example, the most common full-length TprK sequence in AS12 comprised 72.5% of TprK sequences present in that sample and was also present at 3.2% of TprK sequences in MI01. Likewise, a TprK comprising 53.0% of sequences present in MI06 was also present in 0.3% of MI04 TprK sequences.

**Figure 1.**
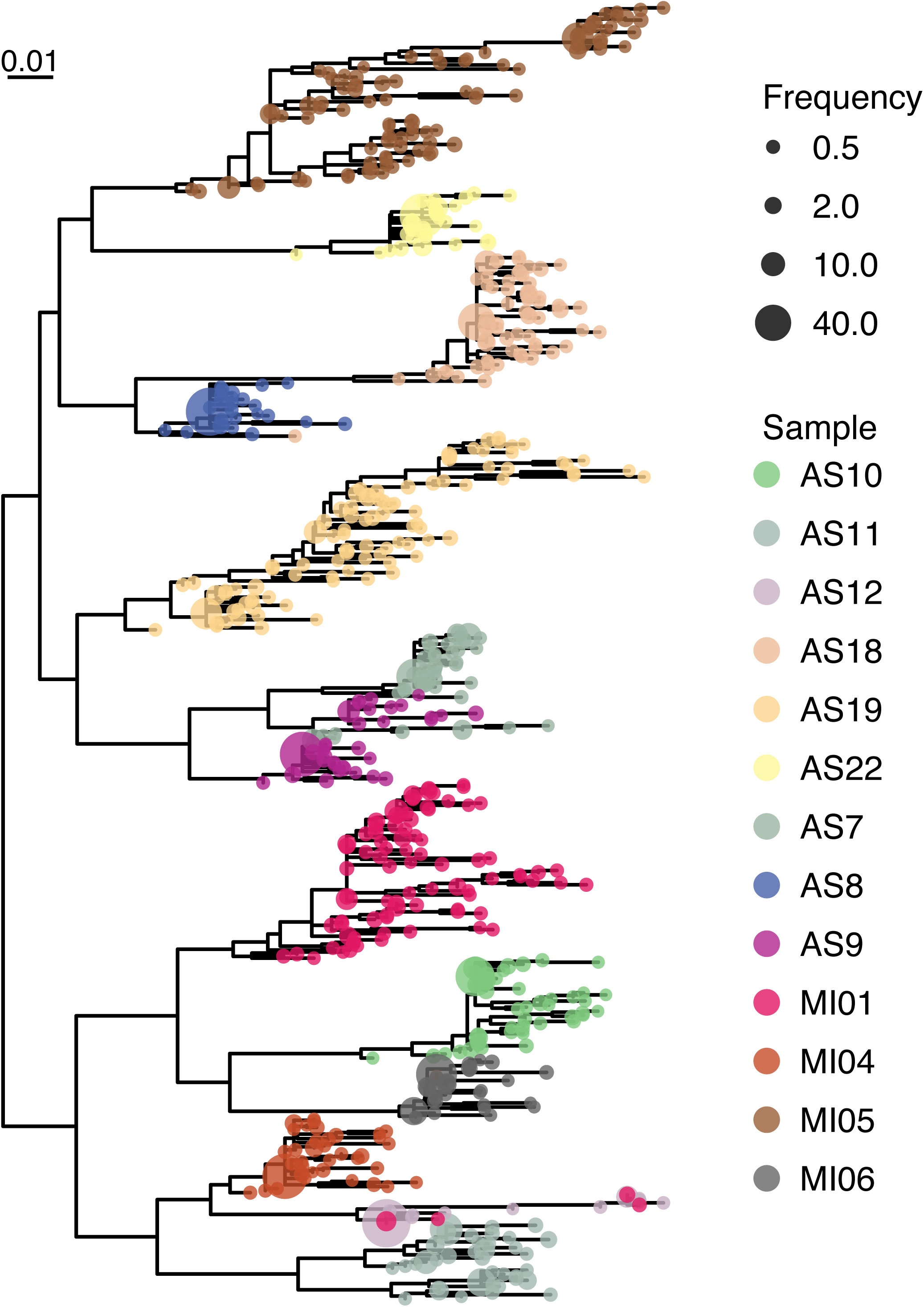
Full-length TprK phylogeny of all protein sequences present at greater than 0.2% within each individual from 13 patients from Italy. Only intact full-length TprK sequences derived from PacBio sequencing were used to generate the phylogenetic tree. Each individual is labeled by a different color and the proportion of sequences is shown by node size. Only three total sequences were shared among the 634 TprK sequences present in the 13 *T*. *pallidum* specimens sequenced in this study.

### Comparison of TprK diversity between Italian and Chinese strains

We next examined whether the TprK V region sequences present in our 13 Italian individuals shared any overlap with TprK sequences derived from short-read sequencing of 28 primary and secondary syphilis specimens recently reported from China [29,30]. Given the extraordinary diversity present in this gene, for print display, we filtered out any variable sequences constituting <20% of the species present in a given sample (Figure 3). More complex data filtered with a minimum frequency of 1% is displayed in an interactive figure in Supplemental File 1.

The heatmap shows the impressive diversity present across the TprK variable regions. V1 and V4 were the most conserved (Figure 2). The same two V1 sequences comprised the highest frequency species present in 9/13 (69.3%) Italian specimens and 16/28 (57.1%) Chinese specimens. Only 12 majority V4 sequences were present across the 41 specimens. However, the most common V4 sequence present in the Chinese samples was only represented once in the Italian cohort and even then it was not the major species present.

**Figure 2.**
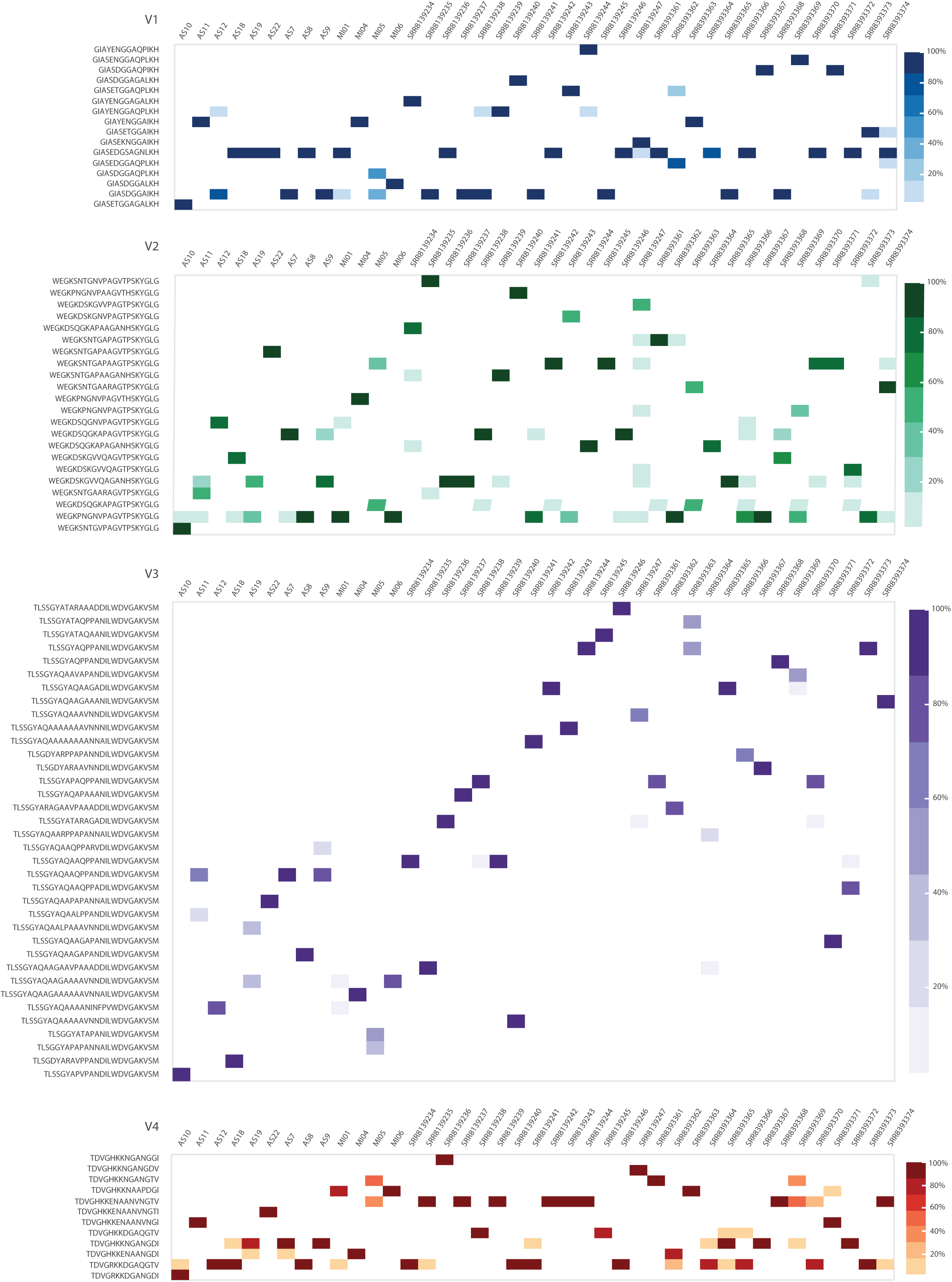
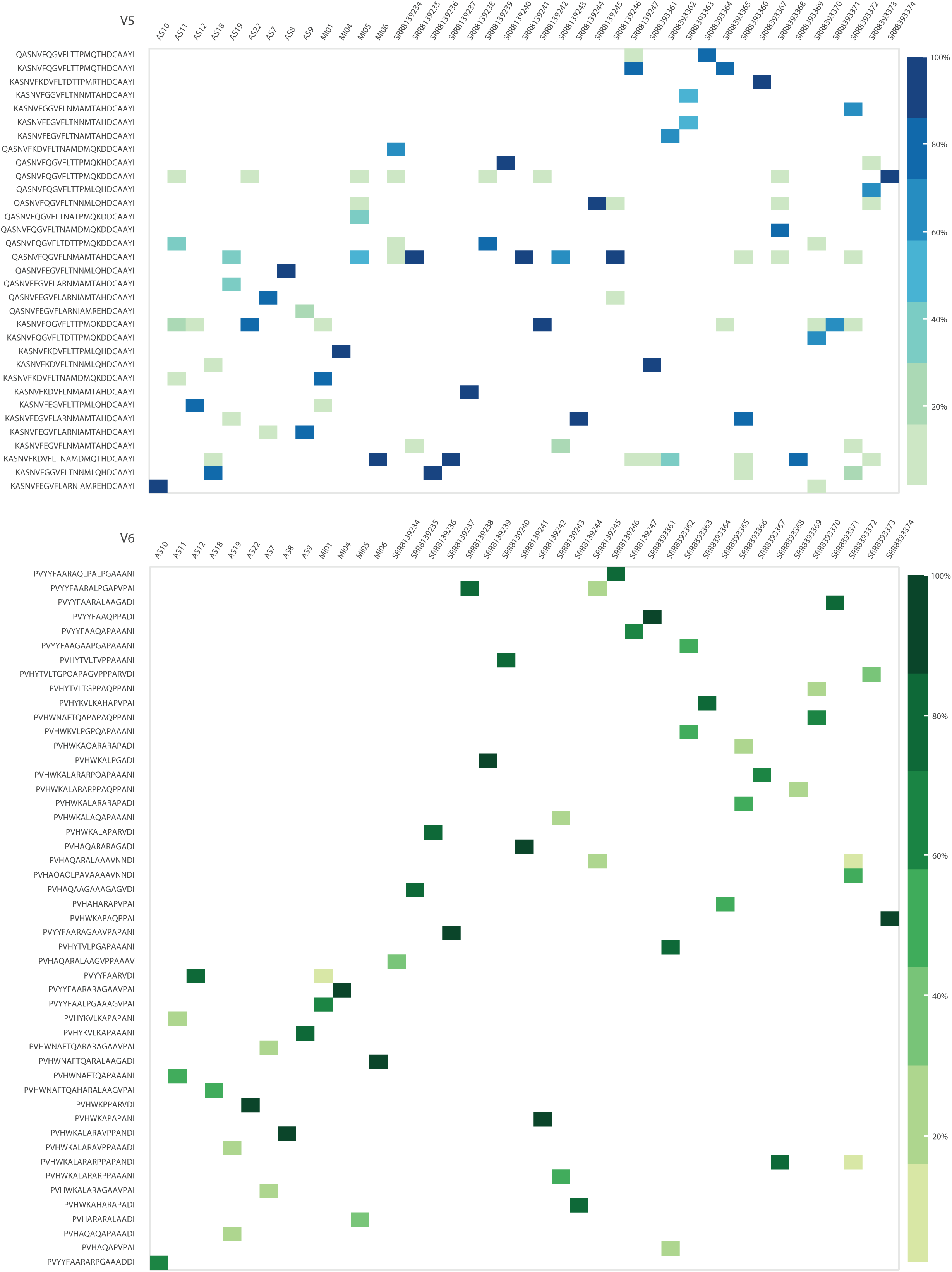
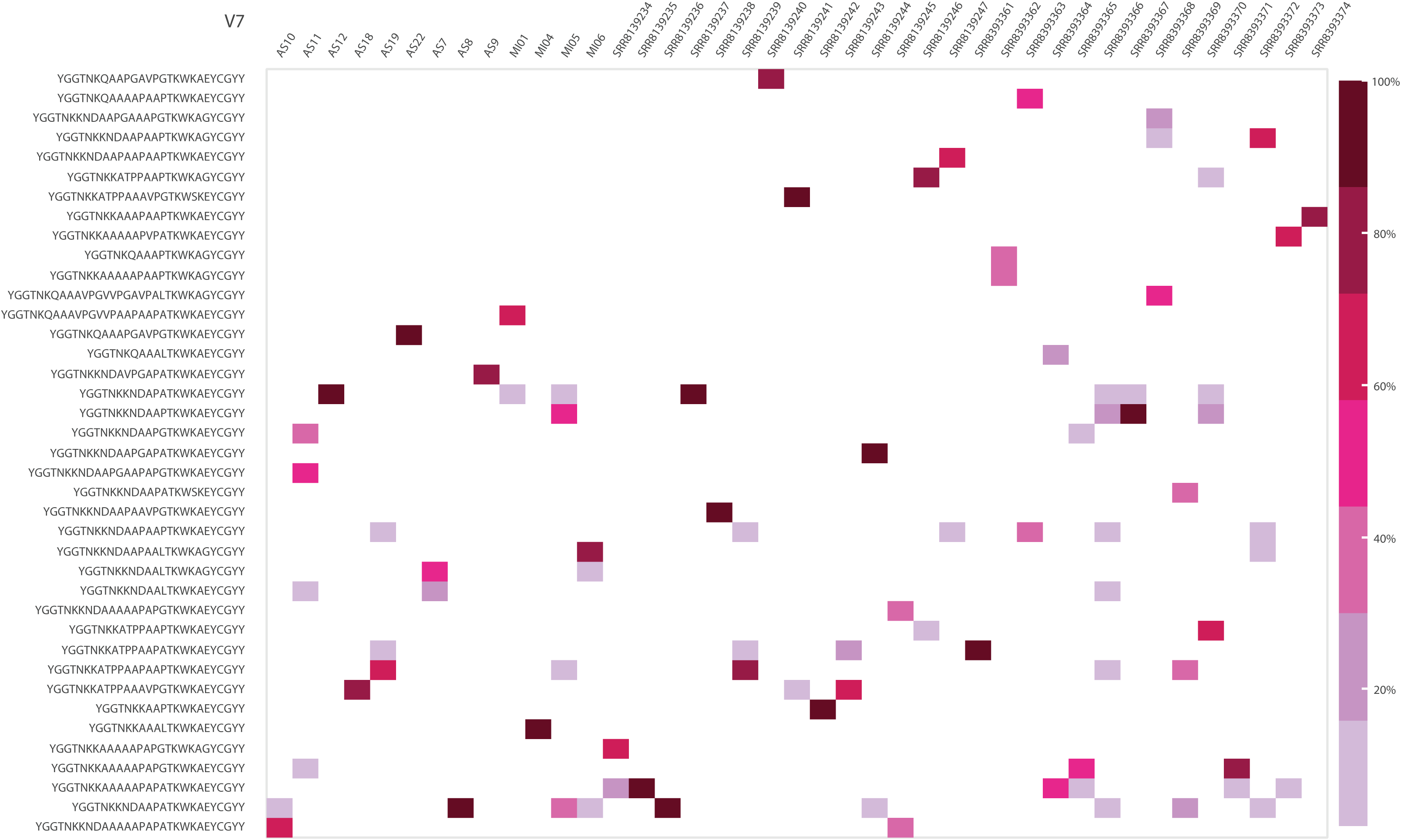
TprK variable region sequence heatmap. Heatmap display of all deep sequenced *tprK* from clinical specimens to date, comprising 13 individuals from Italy sequenced here and 28 Chinese individuals from prior work. For print display, only those variable region sequences present at ≥ 20% frequency within a sample are depicted. Any variable frequencies less than 2% in other samples are not shown. The proportion of sequences is illustrated by color for each heatmap for V1 (A), V2 (B), V3 (C), V4 (D), V5 (E), V6 (F), and V7 (F). A heatmap filtered at a frequency of 1% is provided as an interactive html as Supplemental File 1.

V3, V5, V6, and V7 regions demonstrated almost no overlap among the 41 specimens (Figure 2). Only 6 of 39 major V7 sequences were shared between any Italian and any Chinese specimen. No shared V6 sequences were seen among any samples as the majority species present for each sample.

### Redefining conserved and variable regions in tprK

The sequences we mined from variable regions were initially based off of prior definitions of the conserved and variable portions of *tprK*, which themselves were based off comparatively few *tprK* sequences [25]. While identifying donor sites, we noticed systematic biases in variable region sequence lengths mined from sequencing reads and the total blastn HSP length (Figure S3A/B and reflected in Figure 2). For instance, >98% V3 region sequences started with the same 23 bp sequence (5′-TCATACTCACCTTAGCCCCGACA-3′), and all other sequences had a Levenshtein edit distance of 1 from this sequence, suggesting this sequence may mark a conserved portion of *tprK.* Similarly, 100% of V5 region sequences started with the same 13 bp sequence (5′-AATATAGGCAGCA-3′) and no V5 sequence had less than 13 bp difference in sequence and blast hit length. For V2, 99.3% of sequences began with the same 14 bp sequence (5′-AGTATGGATTGGGG-3′) and the lone alternative sequence could be explained by low-frequency Illumina sequencing error associated with G-quadruplexes [32]. Removal of these sequences improved the ability to align *tprD* donor sites across the length of *tprK* variable region sequences, leaving a 4 nucleotide common sequence (5’-CCTA-3’) in V4 region sequences that we left based on its short nature (Figure S3C/D).

### Contribution of donor sites to variable regions

We next examined how each variable region sequence was generated from different donor sites using data from all 41 samples. We found a total of 55 donor sites, corresponding to 5 for V1, 5 for V2, 13 for V3, 5 for V4, 6 for V5, 14 for V6, and 7 for V7 (Figure 3A). Forty-seven sites were previously reported by Centurion et al. [25]. There was considerable overlap between the two sets, suggesting a finite limit to the number of donor sites for *tprK.* Of note, two donor sites in the *tprD* locus (VS1-15, VS2-21) had single nucleotide variants compared to our reference sequence but exactly matched their previously deposited *tprD* locus (AY587909.1) [25], indicating that chromosomal mutations in donor sites can affect *tprK* variable region sequences. The vast majority of the donor sites found in this analysis, 51/55, were clustered downstream of *tprD*, while the remaining 4 donor sites were located upstream of *tprD*. Notably, all 51 of the donor sites located downstream of *tprD* were in the same orientation as *tprD* and had the highest utilization, while the 4 sites upstream of *tprD* faced in the opposite orientation. Donor sites for specific variable regions were collocated together, such as V1-V4-V5, V2-V7, and V3-V6. V3-V6 donor sites were almost uniformly derived from overlapping sequences (Figure 3B). Donor sites for V1 and V4 were the shortest, measuring an average of 39.2 and 34.8 nucleotides, while V5 and V7 donor sites were the longest at 58.5 and 64.7 nucleotides (Figure 3C).

**Figure 3.**
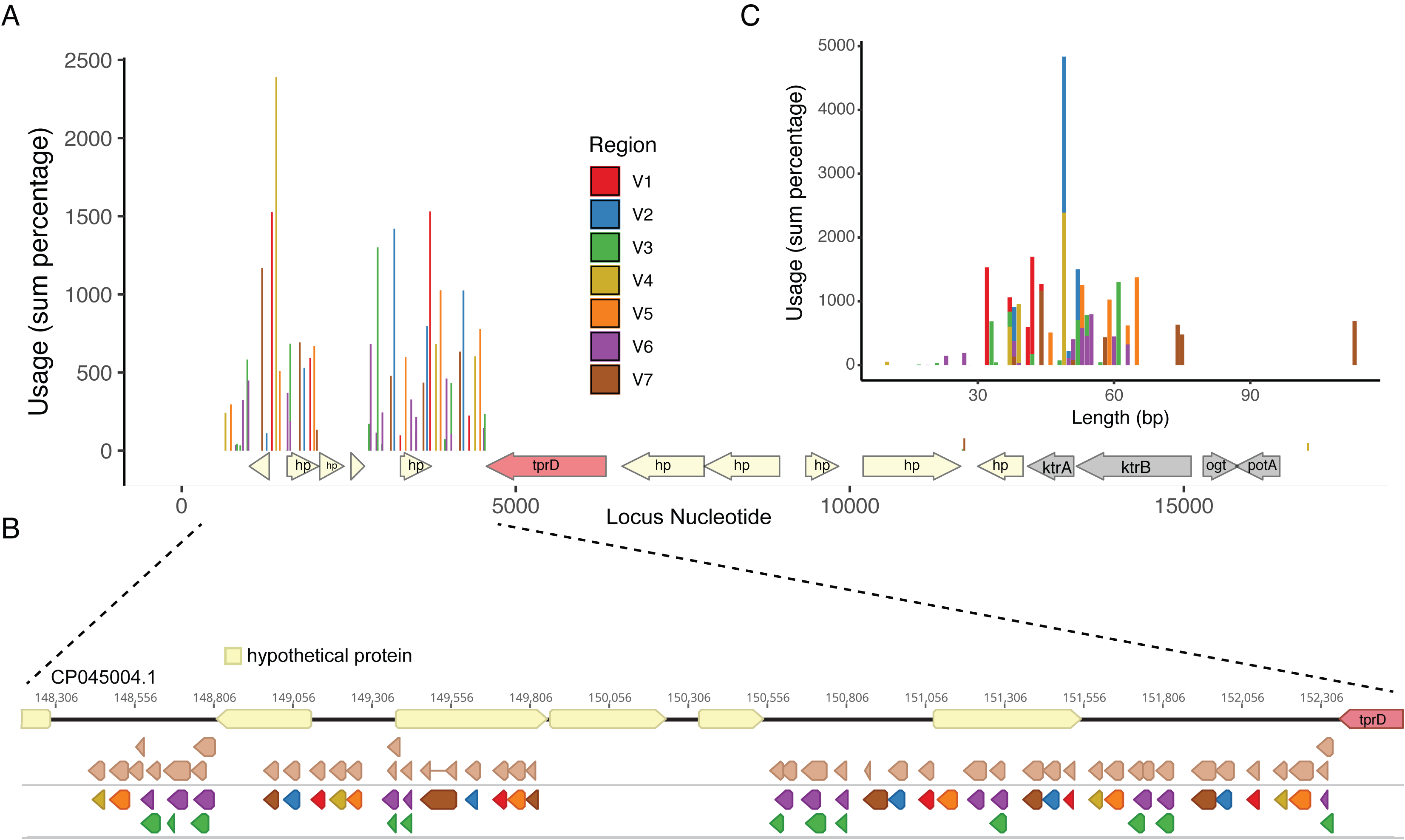
Map of *tprK* donor sites flanking the *tprD* locus. Variable region sequences were blastn aligned against a 17.2kb locus that contained putative *tprK* donor sites based on manual review. A) The usage of all 55 donor sites across the *tprD locus* by variable region is depicted based on the sum of within-sample percentages across all 41 samples. The entire 17.2kb locus is depicted due to recovery of a V4 donor site within the *phnU* gene at 16.9kb. Nucleotide numbering is shown based on the strain UW-148B2 (CP045004.1). B) Zoomed in depiction of the locus immediately downstream of *tprD* containing *tprK* donor sites. Donor sites are in the same orientation as the *tprD* locus. The light brown sites include 45 of the 47 donor sites reported previously by Centurion et al. [25]. The bottom donor sites include 51 of the 55 donor sites found in this study and are colored based on their associated variable region.

### Estimate of total potential diversity of tprK

Using this new inventory of *tprK* donor sites flanking the *tprD* gene, we next estimated the total coding diversity of TprK. Assuming a simple model in which only one donor site contributes to each variable region sequence, the 55 *tprD* donor sites across 7 variable regions could combine to create a total of 955,500 different full-length TprK sequences assuming no mutation. However, multiple donor sites can contribute material to the same *tprK* variable region to create a mosaic variable region. Our manual review of donor site contributions to variable regions suggested that donor sites were limited to three separate contributions to create mosaic variable regions, so we set a limit of three for the number of high scoring pairs in our blastn analysis of donor sites against each variable region sequence. The plurality of V1 region sequences only had one donor site contribute sequence while no V3 or V7 sequences were generated from only one donor site (Figure 4A). However, all variable regions had the potential for three donor site contributions. Adding up all potential combinations of one, two, and three-segment gene conversions that generate different sequences (assuming no single-segment V3 and V7 sequences) and assuming independence between variable regions leads to a potential diversity of TprK of 2.69×10^18^ full-length protein sequences if donor sites are reused, or 1.11×10^17^ protein sequences without reuse (Table S5).

**Figure 4.**
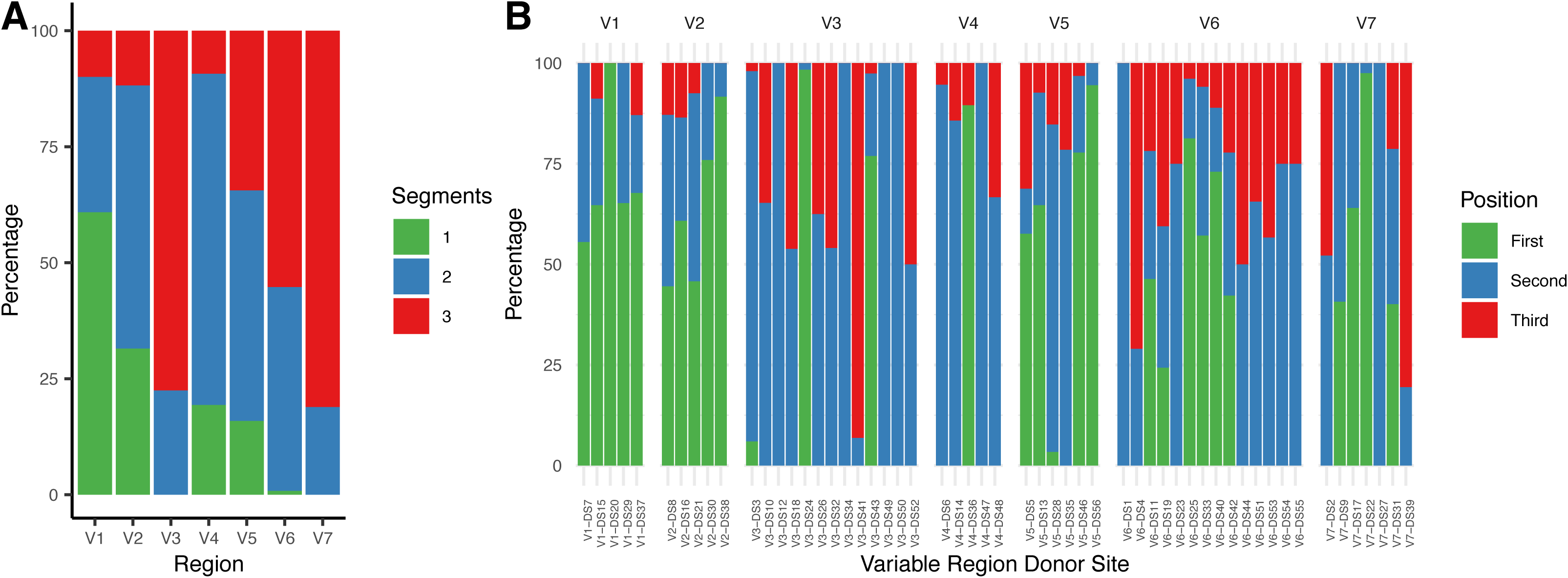
Donor site segments and position by V region. A) The number of donor site contribution segments in each high-confidence variable region sequence was determined in blastn output across the 41 samples. Usage was determined by the sum percentage of variable region sequences by segment. For instance, V1 has the most number of variable region sequences where only one donor site segment is used in a given V region sequence, consistent with its overall lack of diversity. B) The position of donor site contributions within a variable region sequence was also determined for each donor site (i.e. “First” means the donor site was found as the first alignment within a variable region sequence, “Second” as the second portion of the variable region sequence alignment, and “Third” as the third). Within-sample percentages were summed for each variable region in order to adjust for differences in read coverage at each locus between samples. These summed percentages were then adjusted by the total summed percentage to add up to 100% for each variable region.

We next examined whether certain donor sites were not represented in specific sections of a given variable region. Consistent with the segment usage data in Figure 4, we found biases in donor site contribution in every variable region. For instance, every V4 sequence starts with contributions from the same donor site and only two of five total V1 donor sites contribute to the third segment in V1, when using high-confidence variable region sequences (present in more than 50 reads and >0.2% within sample frequency). In addition, V3 and V6 variable regions make use of almost all of their donor sites in both the second and third segments, but less than half of potential donor sites in the first segment. Taking into account differential use of donor sites by variable region segment reduced total potential total diversity to 1.23×10^15^ full-length TprK sequences with replacement, or 7.95 x10^13^ sequences without replacement. Across 1544 individual high-confidence variable region sequences, we found 146 variable region sequences that used the same donor site more than once in the same variable region sequence, indicating that some donor site reuse is allowed in the generation of *tprK* variable regions.

## Discussion

Here we combine deep, full-length profiling of TprK from *T. pallidum*-positive patient specimens with data-mining of additional TprK short-read sequencing from 28 Chinese patients to explore the diversity of the consummate *T. pallidum* immunoevasion protein TprK. We find exceedingly little overlap within specific variable regions within and between each patient cohort. Only 3 of 634 high-quality, full-length TprK sequences were shared among any samples in the 13 patients on which we performed long-read sequencing. Consistent with previous reports, we found greater TprK diversity to be associated with secondary syphilis compared to primary syphilis [29,33]. We then used this dataset of TprK diversity to find additional donor sites and to begin to piece together the grammar of variable region generation.

Based on the lexicon of *tprK* donor sites measured using deep sequencing across 41 samples, we estimate a potential full-length TprK combinatorial diversity approaching 10^14^ – 10^18^ proteins, assuming independence across donor sites. These estimates may be overestimates if our assumption of independence between variable region sequences is incorrect. These estimates may also underestimate the total diversity potential of TprK due to varying lengths of donor site contributions to variable regions. Most importantly, this junctional diversity is similar to if not greater than measures of the human adaptive immune system [34–36].

Our data also provide insights into differences in measured diversity among different variable regions. The limited diversity in V1 is associated with higher use of single-segment gene conversions to generate the variable region, while the limited diversity in V4 is associated with biased positional usage of different donor sites. Using the same or similar numbers of overall donor sites, V2 and V5 are able to generate 2-9 times more possible diversity than V1 and V4, which is reflected in direct sequencing measurements. This increase in diversity generation is due to either less positional bias of donor sites or greater proportions of three-segment donor site contributions, or both.

Of note, we measured fewer than 100 full-length TprKs present in any given sample using our filtering criteria, which is substantially less than our theoretical diversity estimates. Though we demonstrated that increasing our PCR template to the maximum allowed per reaction did not greatly affect variable region diversity measurements, these measured estimates could be biased by the limited copy numbers (<10,000 copies) available for *T. pallidum* positive clinical specimens and the limited range of copy numbers tested in our study.

Our work was chiefly limited by the few numbers of clinical samples and *T. pallidum* strains that have been deeply profiled for TprK diversity. Here, we profiled 13 new *T. pallidum* positive clinical specimens and combined them with 28 previously sequenced samples. However, given the considerable coding potential of TprK, 41 specimens is far too little to understand its overall coding diversity. Because of the limited number of total variable regions sampled across these 41 samples (∼10^3^) versus the potential diversity, we considered ourselves underpowered to examine linkages or epistasis between different variable regions. Future work will have to examine whether certain variable region sequences segregate together within a given TprK. The sampling requirements to determine that association are likely quite considerable and beyond the scope of the work presented here.

In addition to our limited understanding of how TprK variable regions interact with each other, our work here also does not fully inform how TprK interacts with the immune system. As the overall coding diversity of specific variable regions is somewhat limited, it is possible that epistatic interactions between variable regions could influence epitope structure. Certainly, the paucity of variation across the 41 samples in the V4 region is surprising given that anti-V4 antibodies have been detected in humans [37]. We also note the lower number of measured V3 diversity could be associated with a lack of immunological pressure, especially considering its number of potential donor sites and three-segment gene conversions [37]. Also, if there were not epistatic interactions between variable regions, it is not necessarily clear why seven different variable regions with broad but distinct coding potentials would be required in TprK. Alternatively, if there is no or limited epistasis between variable regions and cross-protective antibody is generated against individual variable regions, the diversity generating potential of individual variable regions combined with the rate of gene conversion could put an upward bound on the time period before *T. pallidum* becomes latent in humans.

In summary, our work provides a basis for one mechanism of how *T. pallidum* maintains lifelong infection, through the constant generation of TprK diversity using a lexicon that approaches that of the baseline human adaptive immune system. Therapeutic interventions that target mechanisms of TprK diversity generation may prove beneficial. We further hypothesize that loss of the TprK diversity generation will be one of the first changes associated with longitudinal passage of *T. pallidum* in the new *in vitro* culture system that provides it respite from constant immune selection.

## Supplemental Figure Legends

**Supplemental Figure 1.**
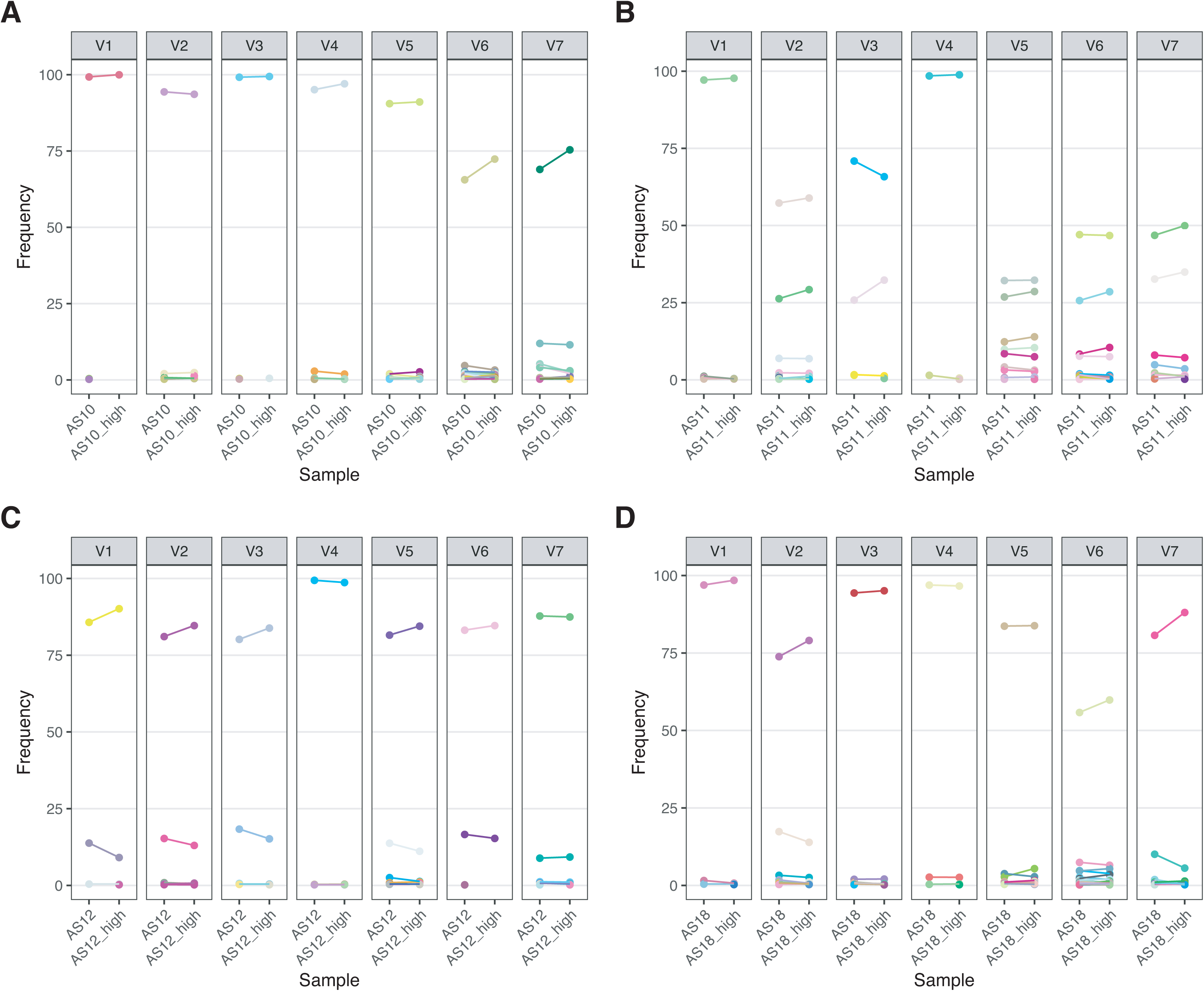
Measurements of diversity are consistent in technical replicates and not significantly affected by template DNA > 1,000 treponemal copies. For strains AS10 (A), AS11 (B), AS12 (C), and AS18 (D) we compared using the maximal template amount allowed by our PCR reaction versus 1,000 treponemal copies. Matched variable regions between the high and normal (1,000 copies) samples are connected by a line and share the same color.

**Supplemental Figure 2.**
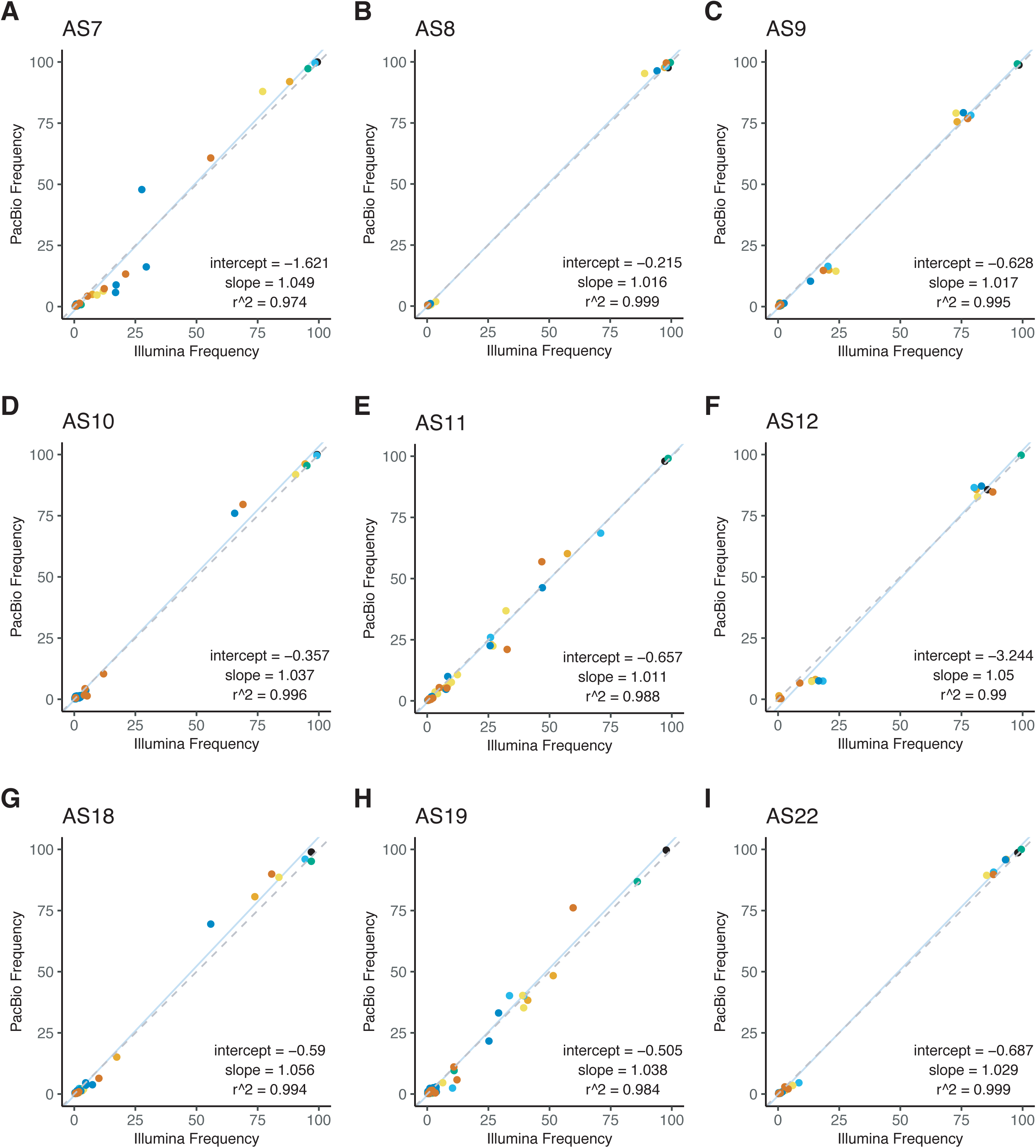
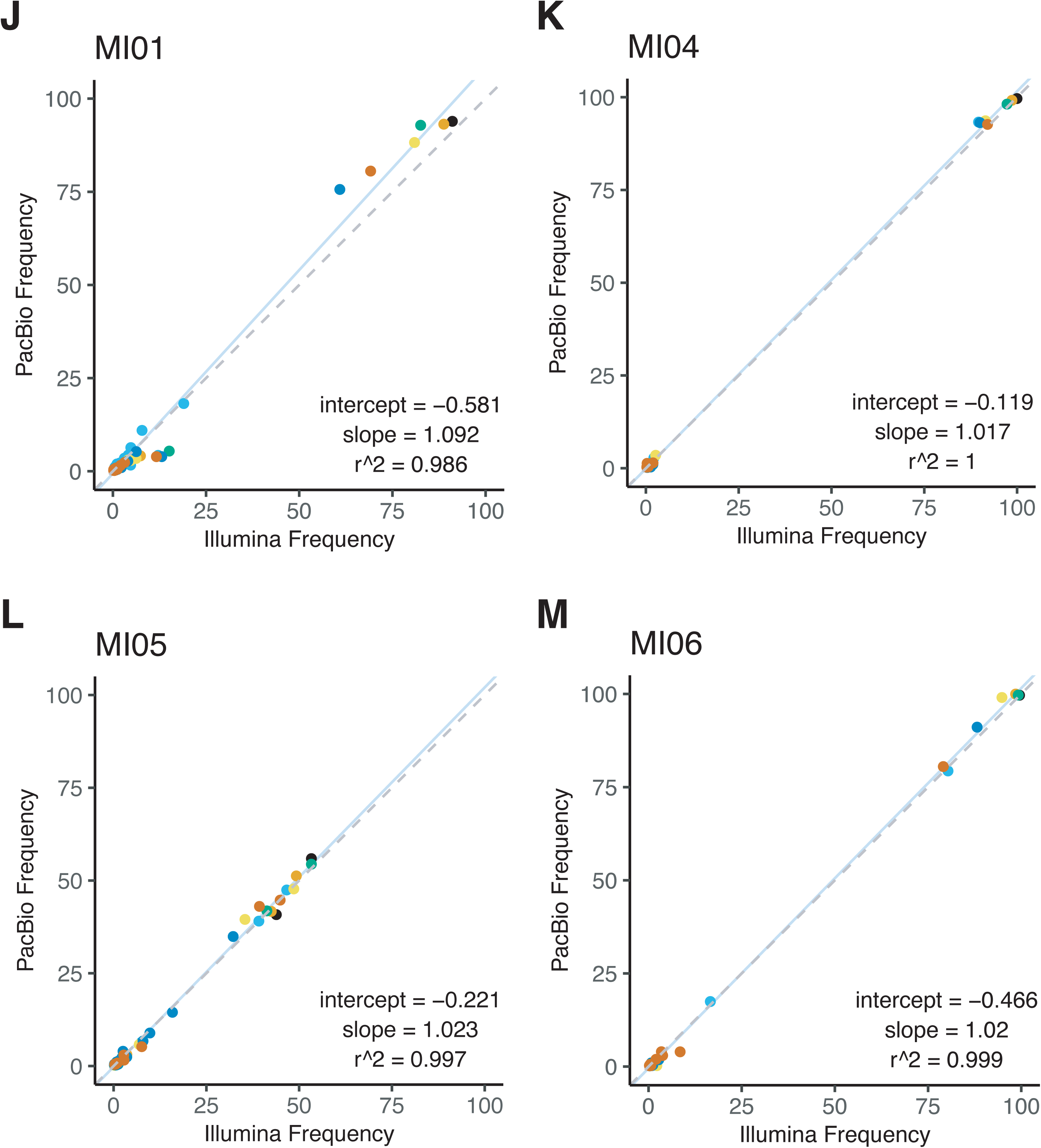
Illumina versus PacBio variable sequence allele frequencies scatterplot for each sample. Each data point is a specific variable region sequence and different regions are labeled by color.

**Supplemental Figure 3.**
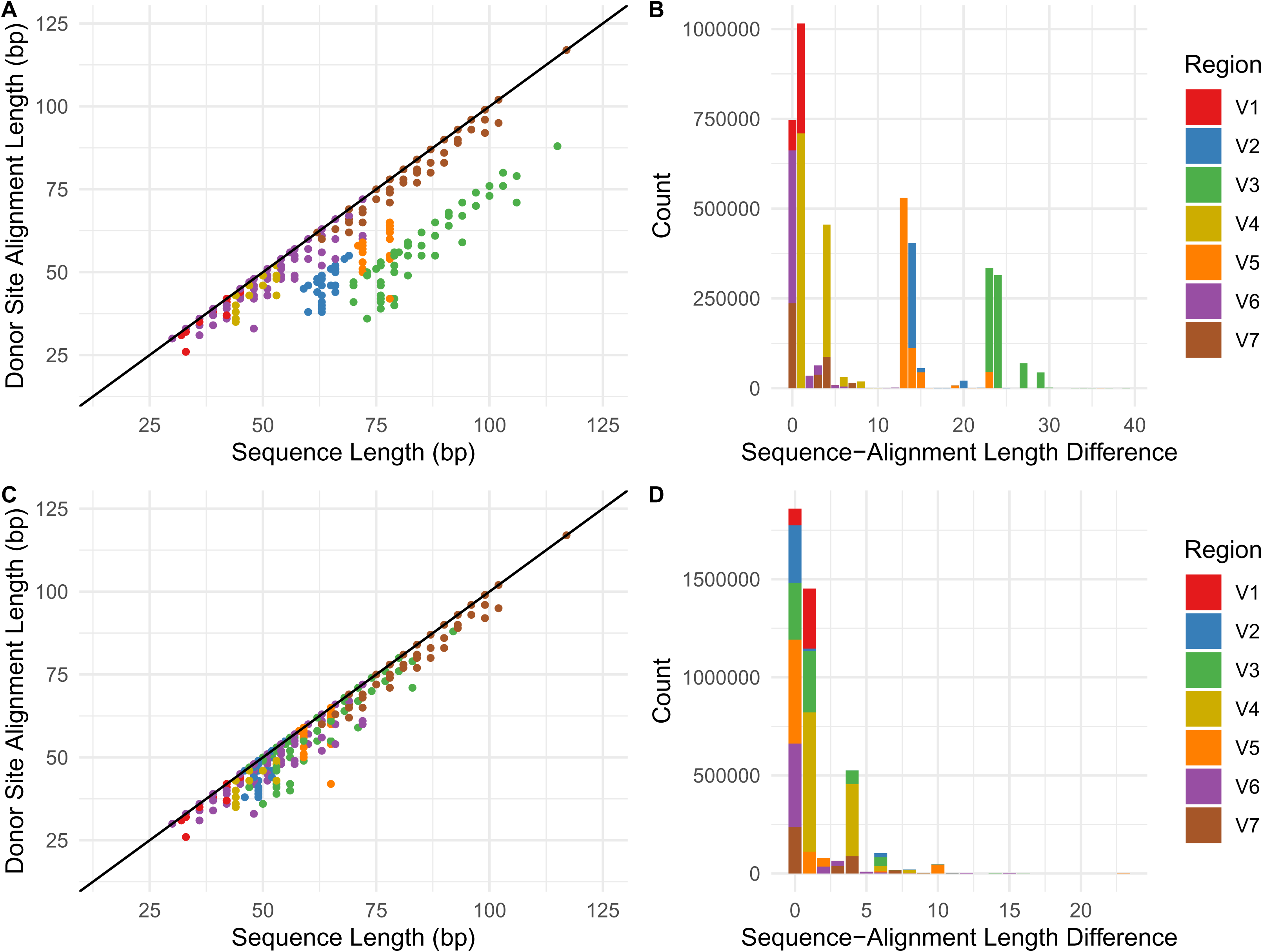
Blast sequence length alignment versus variable region sequence length plots before and after variable region sequence filtering. A) Scatterplot of total sequence length alignment versus variable region sequence length after filtering of V2, V3, and V5 variable region sequences of likely conserved region sequences. B) Corresponding histogram of differences in total sequence length and alignment length after filtering. C) Scatterplot of total sequence length alignment versus variable region sequence length based on prior definitions of variable region sequences. D) Corresponding histogram of differences in total sequence and alignment length without filtering. Counts are absolute sequencing read counts across all 41 samples.

**Supplemental File 1** – Interactive HTML heatmap of TprK V region frequencies across 13 Italian individuals and 28 Chinese individuals. The file contains any variable region sequence present in at least one strain at a frequency greater than 1%.

**Supplemental File 2** – CSV file containing TprK V region sequences extracted from 13 Italian individuals and 28 Chinese individuals previously deep sequenced in *tprK*. Only variable regions present in at least one sample with a minimum of 1% frequency are displayed, as in Supplemental File 1.

**Supplemental File 3** – GFF file of *tprD* locus used in this study with previous donor sites and newly annotated donor sites.

**Table S1.**
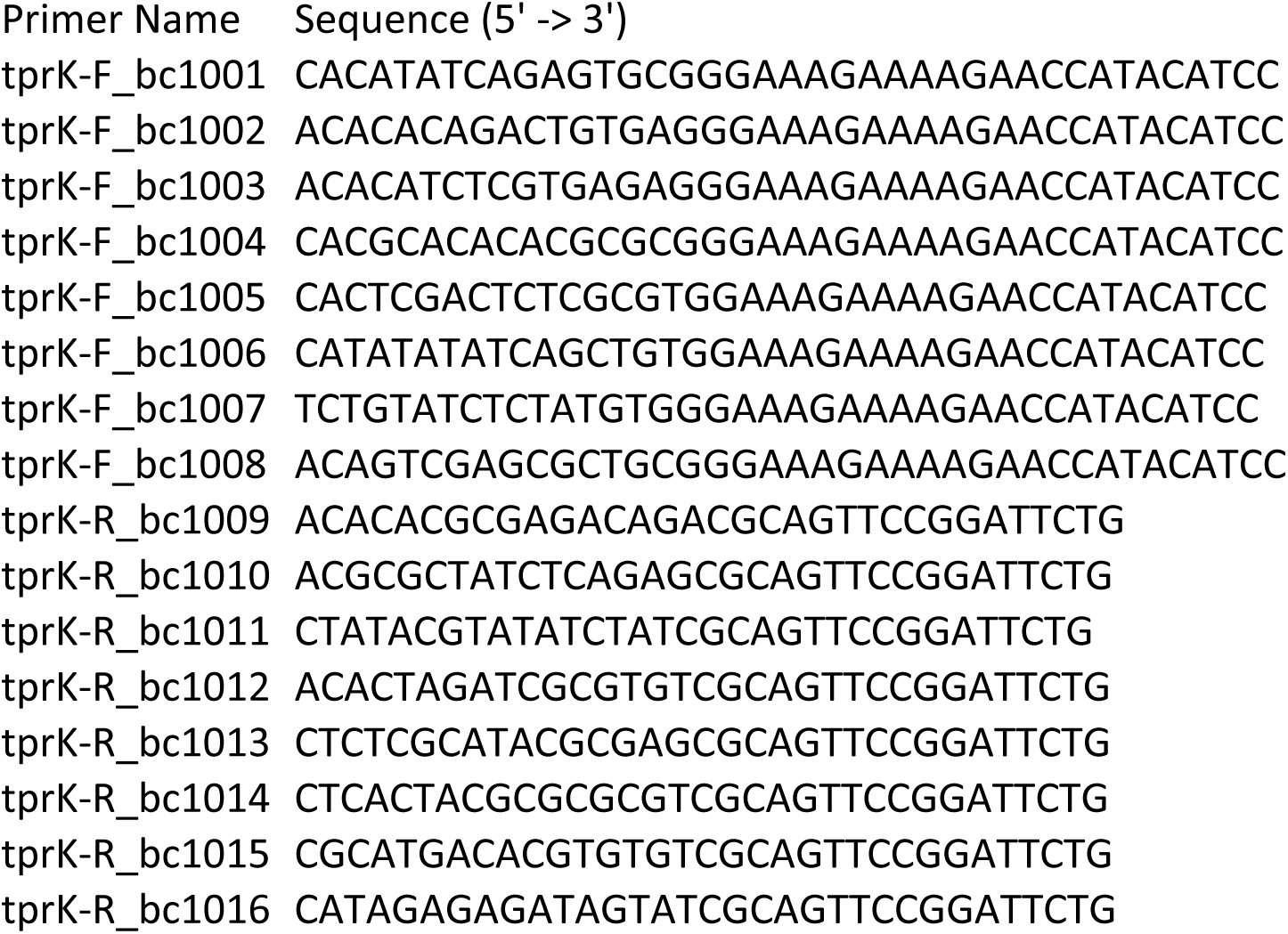
PacBio barcoded *tprK* primers used in this study.

**Table S2.** SRA Accessions for Sequencing Libraries

| SRA Accession | Library |
| --- | --- |
| SRR10953926 | AS7_tprk_nextera |
| SRR10953925 | AS7_tprk_pacbio |
| SRR10953914 | AS8_tprk_nextera |
| SRR10953903 | AS8_tprk_pacbio |
| SRR10953902 | AS9_tprk_nextera |
| SRR10953901 | AS9_tprk_pacbio |
| SRR10953900 | AS10_tprk_nextera |
| SRR10953899 | AS10_tprk_nextera_highcopy |
| SRR10953898 | AS10_tprk_pacbio |
| SRR10953897 | AS11_tprk_nextera |
| SRR10953924 | AS11_tprk_nextera_highcopy |
| SRR10953923 | AS11_tprk_pacbio |
| SRR10953921 | AS12_tprk_nextera_highcopy |
| SRR10953922 | AS12_tprk_nextera |
| SRR10953920 | AS12_tprk_pacbio |
| SRR10953919 | AS18_tprk_nextera |
| SRR10953918 | AS18_tprk_nextera_highcopy |
| SRR10953917 | AS18_tprk_pacbio |
| SRR10953916 | AS19_tprk_nextera |
| SRR10953915 | AS19_tprk_pacbio |
| SRR10953913 | AS22_tprk_nextera |
| SRR10953912 | AS22_tprk_pacbio |
| SRR10953911 | MI01_tprk_nextera |
| SRR10953910 | MI01_tprk_pacbio |
| SRR10953909 | MI04_tprk_nextera |
| SRR10953908 | MI04_tprk_pacbio |
| SRR10953907 | MI05_tprk_nextera |
| SRR10953906 | MI05_tprk_pacbio |
| SRR10953905 | MI06_tprk_nextera |
| SRR10953904 | MI06_tprk_pacbio |

**Table S3.**
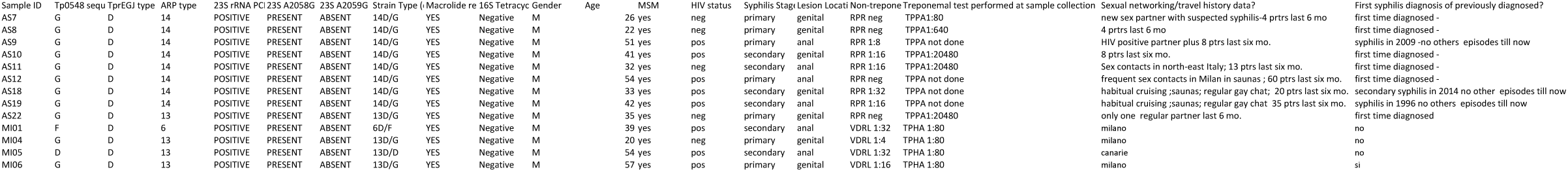
Individual metadata for strains sequenced in this study

**Table S4.**
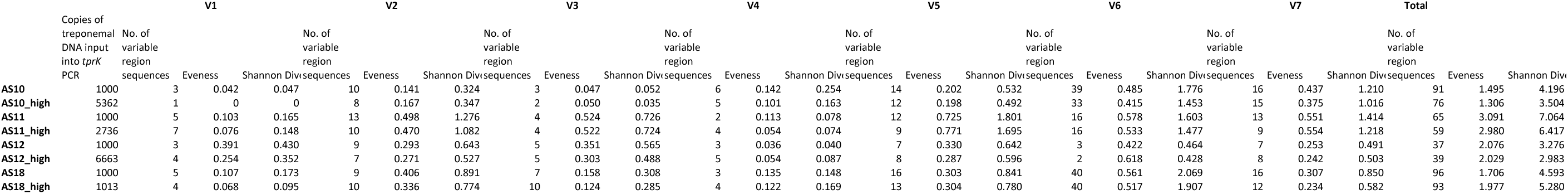
Comparison of the number of variable region sequences and diversity measures identified the 7 variable regions based on t

**Table S5.**
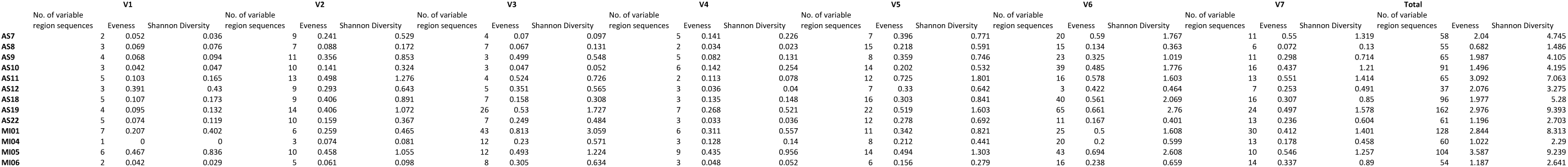
Number of variable region sequences and diversity measures for the 7 variable regions of TprK for the 13 strains profiled in this study.

